# Changes in calpain-2 expression during glioblastoma progression predisposes tumor cells to temozolomide resistance by minimizing DNA damage and p53-dependent apoptosis

**DOI:** 10.1101/2022.10.19.511526

**Authors:** Maren Nicole Stillger, Chia-Yi Chen, Zon Weng Lai, Mujia Li, Agnes Schaefer, Axel Pagenstecher, Christopher Nimsky, Joerg Walter Bartsch, Oliver Schilling

**Affiliations:** Institute for Surgical Pathology, Medical Center – University of Freiburg, Faculty of Medicine, University of Freiburg, Germany; Faculty of Biology, Albert-Ludwigs-University Freiburg, Freiburg, Germany; Internal Medicine Research Unit, Pfizer Inc, Cambridge, MA USA; Department of Pharmaceutical Biology and Biotechnology, Institute of Pharmaceutical Sciences, University of Freiburg, Freiburg, Germany; Department of Neurosurgery, Philipps-University Marburg, Marburg, Germany; Institute of Neuropathology, Philipps-University Marburg, Germany; CMBB, Center for Mind, Brain and Behavior, Marburg University, Hans-Meerwein-Strasse 6, 35032 Marburg, Germany; German Cancer Consortium (DKTK) and German Cancer Research Center (DKFZ), Heidelberg, Germany

**Keywords:** calpain-1, calpain-2, glioblastoma, temozolomide resistance, U251N, DNA damage, TP53

## Abstract

**Background:** Glioblastoma multiforme (GBM) is characterized by an unfavorable prognosis for patients affected. During standard-of-care chemotherapy using temozolomide (TMZ), tumors acquire resistance thereby causing tumor recurrence. Thus, deciphering essential molecular pathways causing TMZ resistance are of high therapeutic relevance.

**Methods:** Mass spectrometry based proteomics were used to study the GBM proteome. Immunohistochemistry staining of human GBM tissue for either calpain-1 or −2 was performed to locate expression of proteases. *In vitro* cell based assays were used to measure cell viability and survival of primary patient-derived GBM cells and established GBM cell lines after TMZ +/− calpain inhibitor administration. shRNA expression knockdowns of either calpain-1 or calpain-2 were generated to study TMZ sensitivity of the specific subunits. The Comet assay and ɣH2AX signal measurements were performed in order to assess the DNA damage amount and recognition. Finally, quantitative real-time PCR of target proteins was applied to differentiate between transcriptional and post-translational regulation.

**Results:** Calcium-dependent calpain proteases, in particular calpain-2, are more abundant in glioblastoma compared to normal brain and increased in patient-matched initial and recurrent glioblastomas. On the cellular level, pharmacological calpain inhibition increased the sensitivities of primary glioblastoma cells towards TMZ. A genetic knockdown of calpain-2 in U251 cells led to increased caspase-3 cleavage and sensitivity to neocarzinostatin, which rapidly induces DNA strand breakage. We hypothesize that calpain-2 causes desensitization of tumor cells against TMZ by preventing strong DNA damage and subsequent apoptosis via post-translational TP53 inhibition. Indeed, proteomic comparison of U251 control vs. U251 calpain-2 knockdown cells highlights perturbed levels of numerous proteins involved in DNA damage response and downstream pathways affecting TP53 and NF-κB signaling. TP53 showed increased protein abundance, but no transcriptional regulation.

**Conclusion:** TMZ-induced cell death in the presence of calpain-2 expression appears to favor DNA repair and promote cell survival. We conclude from our experiments that calpain-2 expression represents a proteomic mode that is associated with higher resistance via “priming” GBM cells to TMZ chemotherapy. Thus, calpain-2 could serve as a prognostic factor for GBM outcome.

## Background

Glioblastoma (GBM), the most frequent brain-derived malignancy, still bears an unfavorable prognosis for patients affected. With a median survival time of 15 months [1], there is an urgent need to develop novel therapies or at least optimize existing ones. Surgical resection followed by radio- and chemotherapy with temozolomide (TMZ) are the standard-of-care treatment modalities [2, 3]. In the course of TMZ therapy, however, tumor and tumor stem-like cells often develop chemoresistance [4, 5], which makes recurrence unavoidable. In this regard, it is crucial to identify essential pathways correlated associated with enhanced TMZ resistance. These pathways can be related to increased capacities in DNA repair, prevention of apoptosis or fundamental changes in cell metabolism. Generally, proteins increased under therapy of glioblastoma are potentially able to desensitize glioblastoma cells against adjuvant therapy regimens. As such proteins, the calcium-dependent proteases calpain-1 (CAPN1) and −2 (CAPN2) as well as their small regulatory subunit CAPNS1 were identified based on their increased abundance in the present study.

Calpains are non-lysosomal, cysteine proteases, and around 15 human calpain proteases have been discovered so far [6]. As calcium-dependent proteases, calpains require different calcium concentrations for their activation *in vitro* [7, 8]. While calpain-1 and −2 are ubiquitously expressed, the other family members show tissue-specific expression patterns (reviewed in [6, 9]). Both main calpain proteases form heterodimers with the shared, small regulatory subunit CAPNS1 [10]. Further, calpain activity is regulated by its endogenous inhibitor calpastatin which binds up to four calpain heterodimers simultaneously [11–13]. With regard to their cleavage specificity, calpain proteases do not strictly recognize specific amino acid sequences, but show a preference for higher-order structures [14, 15]. Instead of merely proteolytically degrading their substrates, they often modulate protein activity in cell motility, cell adhesion, autophagy and apoptosis (reviewed in [6, 9]). Moreover, calpain proteases are involved in various disorders such as Alzheimer’s disease [16–18], Parkinson’s disease [19], glioma [20], colorectal cancer [21, 22], acute myeloid leukaemia [23], or breast cancer [24, 25]. In a recent study, calpain-1 activity was used to stain glioblastoma during surgical intervention leading to a spatial distinction between tumor and peritumoral tissue [26]. While the implication of calpain proteases in cytoskeletal remodeling, cellular migration, and invasion in cancer biology is well-established, their role in apoptosis is rather unclear [9]. Pro-survival functions link calpains to the degradation of the tumor-suppressor protein p53 (TP53) [27–32], IкBα [33, 34], the inhibitor of NF-кB, and the transcription factor MYC [35]. Pro-apoptotic functions imply the cleavage and thereby activation of caspase-7, −10 and −12 [36–39]. Further, apoptosis promoting functions involve the cleavage of pro-apoptotic proteins including BAX [40, 41], CDK5 [42], APAF1 [43], JNK [39], JUN or FOS [44, 45]. However, most of these studies were conducted in models of neuronal cell death rather than in cancer models showing the necessity to study the role of calpains in cancer biology.

In the present study, we used patient samples of initial and recurrent glioblastoma as well as of non-malignant brain tissue to study the expression profiles of calpain-1 and calpain-2. Furthermore, we examined the role of calpain proteases in TMZ resistance using primary and established glioblastoma cell lines. We observed a strong overexpression of calpain-2 in glioblastoma on both, protein and mRNA levels. Moreover, we show that calpain inhibition in primary cells as well as in an established glioblastoma cell line (U251N) increases their sensitivity to TMZ. Functionally, we show that knockdown of calpain-2 results in increased caspase-3 cleavage and the dysregulation of DNA damage sensing and repair proteins before any treatment with DNA damage-inducing agents. Finally, we link the pro-apoptotic impact of silenced calpain-2 expression to an increased TP53 protein abundance.

## Material and Methods

### Antibodies

Antibodies directed against calpain-1 (ab39170), calpain-2 (ab39165), GAPDH (ab8245) and tubulin beta III (ab7751) were obtained from Abcam; Calpastatin from Santa Cruz (sc376547), cleaved caspase-3 (Asp175) (5A1E) (#9664) and phosphor-histone H2A.X (Ser139) (#2577) from Cell Signaling Technology, TP53 (21891-1-AP) from Proteintech, and 53BP1 clone BP13 (MAB3802) from Sigma-Aldrich. Secondary horseradish peroxidase (HRP) conjugated anti-mouse (#1721011) and anti-rabbit (#1721019) antibodies were obtained from BIO-RAD Laboratories. The AF555 anti-rabbit (ab150078) and AF647 anti-mouse (ab150115) antibodies were obtained from Abcam.

### Inhibitors and drug administration

Temozolomide (TMZ, T2577) and neocarzinostatin (NCS, N9162) were both purchased from Sigma-Aldrich. Calpains were inhibited using 50 µM PD150606 (Calbiochem, 513022). 20 µM Q-VD-OPh (Sigma-Aldrich, SML0063) were used to inhibit caspases; 100 µM Nec-1 (Sigma-Aldrich, N9037) to inhibit necroptosis. We used 100 nM of the BH3 mimetic ABT-737 (Selleckchem) and 1 µM of the Mcl-1 inhibitor S63845 (Selleckchem) to induce apoptosis. Except for NCS, all chemicals were dissolved in DMSO (A3672, AppliChem). NCS is delivered in 20 mM 2-(N-morpholino)ethanesulfonic acid (MES) buffer pH 5.5 by the manufacturer.

### Ethics

In accordance with the local ethics committee of the Medical Faculty, Philipps University Marburg, ethical approval was obtained under file number 185/11. Tumor tissue samples of GBM patients were obtained during surgical resection and all patients provided written informed consent prior to tumor resection.

### Patient samples

Tissue samples were shock frozen in liquid nitrogen during surgery and then stored at −80 °C. Non-malignant brain tissue was collected when tumor sites were accessed during neurosurgery from non-matched patients. All included brain tumor tissues were isocitrate dehydrogenase (IDH) wild type GBM tumors and classified according to the new WHO classification scheme from 2021 as CNS WHO grade 4. In 19 cases, tumor materials from recurrent tumors were available, so that matched pairs of initial (iGBM) and recurrent (rGBM) glioblastoma were studied. All relevant clinical information and histopathological characteristics are summarized in Table 1 and Table 2.

**Table 1:**
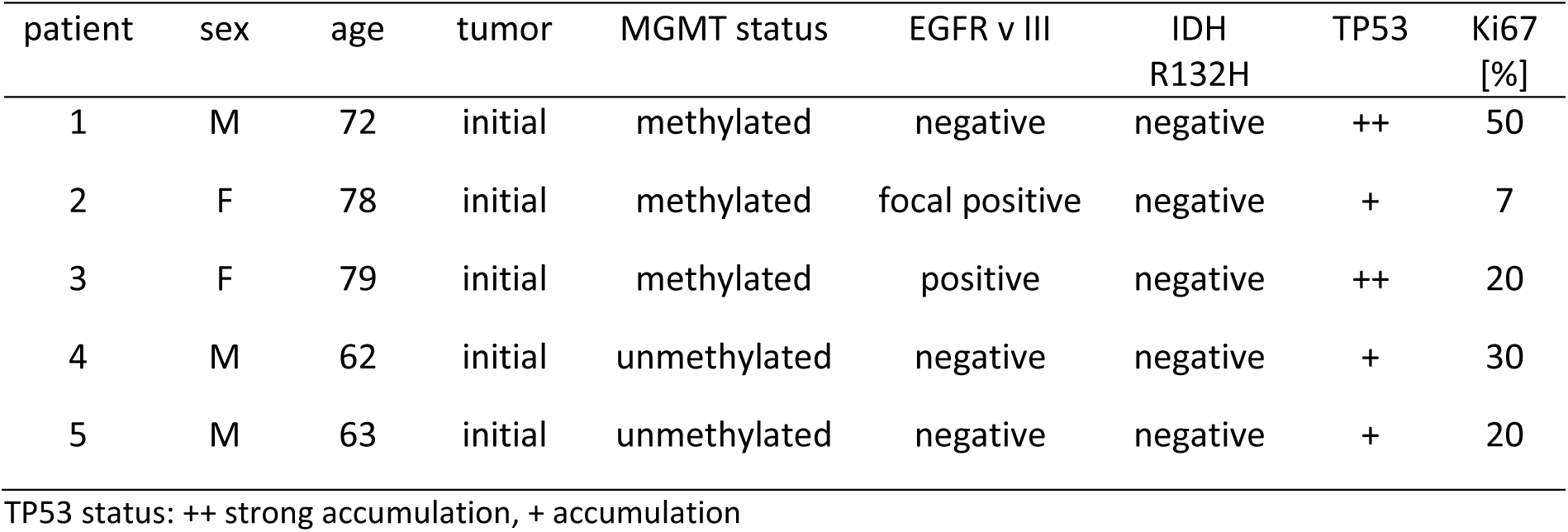
Clinical data of patients included in the proteome comparison of GBM vs. non-malignant brain (resected from the same patient to approach the tumor site during surgery).

**Table 2:** Clinical data for GBM patients included in the calpain mRNA expression profiling.

| patient | sex | survival<br>[days] | tumor | MGMT status | EGFR v III | IDH r132H | Tp53 | Ki67<br>[%] |
| --- | --- | --- | --- | --- | --- | --- | --- | --- |
| 1 | m | 511 | initial | unmethylated | negative | negative | 0 | 25 |
|  |  |  | recurrent | unmethylated | negative | negative | ++ | 10 |
| 2 | m | 553 | initial | unmethylated | negative | negative | ++ | 20 |
|  |  |  | recurrent | unmethylated | negative | negative | ++ | 30 |
| 3 | m | 543 | initial | unmethylated | negative | negative | - | 30 |
|  |  |  | recurrent | unmethylated | negative | negative | - | 30 |
| 4 | m | 371 | initial | unmethylated | negative | negative | - | 25 |
|  |  |  | recurrent | unmethylated | negative | negative | + | 10 |
| 5 | m | 387 | initial | unmethylated | negative | negative | 0 | 50 |
|  |  |  | recurrent | unmethylated | negative | negative | + | 25 |
| 6 | m | 760 | initial | unmethylated | focal exp. | negative | focal + | 75 |
|  |  |  | recurrent | unmethylated | negative | negative | focal + | 20 |
| 7 | f | 445 | initial | unmethylated | negative | negative | partial + | 20 |
|  |  |  | recurrent | unmethylated | negative | negative | + | 5 |
| 8 | m | 343 | initial | unmethylated | negative | negative | + | 50 |
|  |  |  | recurrent | unmethylated | negative | negative | ++ | 10 |
| 9 | f | 550 | initial | unmethylated | negative | negative | ++ | 15 |
|  |  |  | recurrent | unmethylated | focal exp. | negative | 0 | 7 |
| 10 | m | 524 | initial | unmethylated | negative | negative | 0 | 25 |
|  |  |  | recurrent | unmethylated | negative | negative | ++ | 5 |
| 11 | f | 1277 | initial | methylated | negative | negative | ++ | 50 |
|  |  |  | recurrent | methylated | negative | negative | + | 40 |
| 12 | f | 494 | initial | unmethylated | negative | negative | ++ | 10 |
|  |  |  | recurrent | unmethylated | negative | negative | ++ | 20 |
| 13 | m | 576 | initial | unmethylated | negative | negative | - | 30 |
|  |  |  | recurrent | unmethylated | negative | negative | + | 5 |
| 14 | f | 1997 | initial | unmethylated | negative | negative | + | 10 |
|  |  |  | recurrent | unmethylated | negative | negative | + | 20 |
| 15 | f | 679 | initial | unmethylated | negative | negative | 0 | 40 |
|  |  |  | recurrent | unmethylated | negative | negative | + | 50 |
| 16 | f | 869 | initial | methylated | negative | negative | + | 30 |
|  |  |  | recurrent | methylated | negative | negative | + | 5 |
| 17 | m | 457 | initial | unmethylated | negative | negative | ++ | 20 |
|  |  |  | recurrent | unmethylated | negative | negative | - | 3 |
| 18 | f | 328 | initial | methylated | negative | negative | ++ | 20 |
|  |  |  | recurrent | methylated | negative | negative | + | 10 |
| 19 | m | 1624 | initial | unmethylated | negative | negative | ++ | 40 |
|  |  |  | recurrent | unmethylated | negative | negative | ++ | 5 |

TP53 status: ++ strong accumulation, + accumulation, 0 moderate accumulation, - weak accumulation

### Explorative Mass Spectrometry (MS)-based Proteomics of iGBM and non-malignant brain tissue

Proteins from fresh frozen human brain tissue specimens were extracted and prepared as previously described [46, 47]. In brief, one initial GBM and one non-malignant brain sample were arranged as pairs and labelled with stable isotope variants of formaldehyde as described previously [47]. Peptide concentrations were determined with the Pierce^TM^ BCA Protein Assay Kit according to the manufacturer’s instructions (#23225, ThermoFisher). Samples were measured on an Orbitrap Q-Exactive plus (Thermo Scientific^TM^) mass spectrometer coupled to an Easy nanoLC 1000 (Thermo ScientificTM) as described previously [47]. Data were analyzed using MaxQuant Version 1.5.2.8 [48] with the human reference database downloaded from UniProt on the 15^th^ of April 2016 (20,193 protein entries). Missed tryptic cleavages were not allowed, first search tolerance was set to 20 ppm, main search tolerance to 4.5 ppm. Carbamidomethylation of cysteine was set as a fixed modification. No variable modifications were set. The false discovery rate for peptides and proteins was set to 0.01.

### Isolation of primary cells from GBM patients

Primary patient-derived GBM cells were prepared from CNS WHO grade 4 specimens collected directly after surgery as described earlier [49]. Tumor tissues were washed in HEPES-buffered saline, homogenized and treated for 30 min with 0.025 % Trypsin/EDTA solution at 37 °C. The resulting cell homogenate was passed over an 80 µm cell strainer and the cell suspension was centrifuged (200g, 5 min). After two washes with medium (DMEM, 10 % FCS), the cells were seeded out for propagation and kept under differentiating conditions.

### GBM cell culture

A172, U87, T98G, U251(N) and primary GBM cells were cultured in Dulbecco’s modified Eagle’s medium (DMEM) high glucose (Gibco^TM^, ThermoFisher) with 10 % FBS (P30-3031, Pan Biotech) and 1 % penicillin-streptomycin (Gibco^TM^, ThermoFisher) in a humidified incubator at 37 °C with 5 % CO_2_. Depending on confluency, cells were subcultured every 3-4 days by washing with Dulbecco’s phosphate buffered saline (DPBS, Gibco^TM^, ThermoFisher) and detached with trypsin (Gibco^TM^, ThermoFisher). Cell line identity was authenticated using the short tandem repeat loci (STRs) method by Microsynth AG (Switzerland). Cell cultures were tested regularly for mycoplasma contamination by eurofins Genomics (Germany).

### Immunohistochemical staining for CAPN1 and CAPN2

Formalin fixed and paraffin embedded tissue sections (3 µm) were stained using the VECTA Stain Elite Kit (Vecta Laboratories, Burlingame, USA) according to manufacturer’s instructions. After deparaffinization at 60 °C for 45 min, sections were rehydrated using descending alcohol concentrations. Demasking of epitopes was achieved by boiling in citrate buffer (10 mM trisodium citrate dihydrate, pH 6). Endogenous peroxidase was blocked by incubating in 3 % H2O2 in methanol for 30 min before incubation in 1.5 % goat serum for blocking of unspecific binding. Followed by incubation with primary antibody (anti-CAPN1, Abcam ab39170, 1:200; anti-CAPN2, Abcam ab39165, 1:100; diluted in PBS) at 4 °C overnight. Incubation with the respective secondary biotinylated antibody and ABC reagent were followed by DAB staining with the ImmPACTT DAB Kit (Vecta Laboratories, Burlingame, USA). For counterstaining, hematoxylin (Carl Roth, Karlsruhe, Germany) was used. Finally, sections were dehydrated by ascending ethanol concentrations and covered with mounting medium. Images were acquired using a Leica Axiophot XX with an integrated camera system.

### Real-time quantitative polymerase chain reaction

To extract RNA, 50 mg tumor tissue was mechanically homogenized in 1 ml Qiazol (Qiagen, Hilden, Germany), whereas cells were lysed in 1 ml Qiazol by resuspending. 200 µl of chloroform were added, samples were mixed and incubated for 3 min followed by centrifugation for 15 min at 4 °C. The upper aqueous phase was added to 500 µl Isopropanol to precipitate RNA. The pellet was washed with 75 % ethanol, dried and solved in RNase free water. For reverse transcription 2 µg RNA were translated into cDNA using the “RNA to cDNA EcoDry” Kit (Takara Bio USA, Kusatsu, Japan) according to manufacturer’s instructions. The following qPCR analyses were performed in triplicates using the “Precision FAST MasterMix with ROX” (Primer Design, Southampton, UK), the respective Quantitect Primer pairs for detection of specific mRNAs (Qiagen, Hilden, Germany), and StepOnePlusTM qPCR system (2^-ΔΔCT^ method). Values were normalized to endogenous housekeeping genes (RPL7 or RPLP0).

### Cell viability assays

To measure ATP consumption, a CellTiter-Glo® luminescent cell viability assay (G7570, Promega) was used. Cells were seeded on 96-well plates (1000 cells/well, triplicates) and treated with specified conditions. After 5-day treatment, cell viability was examined according to the manufacturer’s instructions. In brief, the CellTiter-Glo® solution was applied to each well and incubated at room temperature for 10 min. The intensity of luminescence was measured using a microplate reader (EnSpire, PerkinElemer). Cell viability was calculated relative to DMSO treatment (100 %). For the MTT proliferation assay, cells were seeded on 96-well plates (5000 cells/well, triplicates) and cultured under indicated conditions. For each condition, the reagent 3-(4,5-Dimethylthiazol-2-yl)-2,5-diphenyltetrazolium bromide (MTT, Sigma-Aldrich) was added to the cells and incubated for 4 hours at 37 °C (5 % CO_2_). The supernatants were discarded and replaced by 200 µL DMSO to dissolve the formazan product. Absorbance was measured at 570 nm in a plate reader (BMG Labtech).

### Propidium iodide and Hoechst 33342 staining for dead rate counting

Cells were seeded on 24-well plates (5000 cells/well, triplicates) and treated with the specified condition. After 3- or 5-day treatment, cells were washed with DPBS and staining solutions were applied according to the manufacturer’s instructions (ReadyProbes® Cell Viability Imaging Kit, R37610, ThermoFisher). Cells were incubated at 37 °C for 15 min. Blue (Hoechst 33342) and red (propidium iodide) staining cells were inspected by microscopy (Axio Observer Z1 Zeiss). The live/dead cell number was counted using Fiji ImageJ [50].

### Cell counting

Cells were seeded on 10 cm dishes (2.5*10^4^ cells/dish, triplicates) and treated with the specified condition. After 24 hours, cells were washed with DPBS and detached with trypsin. Cells were stained with trypan blue to identify dead cells and counted with the EVE^TM^ automatic cell counter system (NanoEnTek).

### Calpain activity assay

Cells were seeded on 10 cm dishes (10^5^ cells/dish, triplicates) and treated with the specified condition. After 5-day treatment, calpain activity assay was performed according to the manufacturer’s instructions (ab65308, Abcam). In brief, cells were washed with cold DPBS and lysed with the extraction buffer. To increase the extraction efficiency, cells were homogenised by going 10 times through needles (27 G, B. Braun Melsungen AG). After incubation on ice for 10 min, supernatant was collected by centrifugation 10 min, 13,500 rpm at 4 °C. 25 μg of protein was incubated with the calpain substrates at 37 °C for 1 h. The fluorometric intensity (Ex/Em 400/505 nm) was measured using a microplate reader (EnSpire, PerkinElemer). Calpain activity was calculated relative to DMSO treatment for each cell line.

### CAPN1 and CAPN2 expression knockdown in U251N cell lines

Expression of either CAPN1 or CAPN2 was knocked down in U251N using shRNA purchased from Sigma-Aldrich (CAPN1: TRCN0000432907, TRCN0000431534; and CAPN2: TRCN0000003540, TRCN0000284853; non-target shRNA Control Plasmid: SHC016). We used a lentiviral packaging system (SHP001, Sigma-Aldrich) together with lipofectamine™ 3000 (L3000015, Invitrogen™ ThermoFisher) transfection to produce virus in HEK293 cells. Crude supernatants containing viruses were applied onto U251N cells to transduce those cells and to induce stable expression knockdowns. Expression knockdowns were verified by western blotting.

### Western Blotting

Cells were seeded on 10 cm dishes and (10^5^ cells/plate, triplicates) and treated with the specified condition. After treatment, cells were washed with ice cold DPBS. 500 µl RIPA buffer (150 mM sodium chloride, 1 % Triton X-100, 0.5 % sodium deoxycholate, 0.1 % sodium dodecyl sulfate, 50 mM Tris, pH 8.0) were applied and incubated on ice for 10 min. Protein extracts were collected with a cell scraper and centrifuged for 10 min at 13,500 rpm to remove cell debris. Protein concentration was measured with the Pierce^TM^ BCA Protein Assay Kit – Reducing Agent Compatible according to the manufacturer’s instructions (#23250, ThermoFisher). 30 µg total protein extract was mixed with 4x Laemmli Sample Buffer (#1610747, BIORAD) and denatured for 10 min at 95 °C. After electrophoresis, proteins were transferred onto a 40 µm PVDF membrane (88518, Thermo Scientific^TM^). Membranes were blocked with 5 % bovine serum albumin (BSA, Sigma-Aldrich). Target proteins were detected with specific antibodies. HRP conjugated antibodies were diluted 1:5,000. Chemiluminescence substrate (SuperSignal^TM^ West Femto Maximum Sensitivity Substrate, 34094, Thermo Scientific^TM^) was applied to detect the chemiluminescent signal with the Fusion FX (Vilber Lourmat).

### Explorative MS-based proteomics of U251N shCTRL and shCAPN2

To compare the proteomes of the U251N shCTRL and shCAPN2 cells, 10^6^ cells were collected and washed with DPBS. Proteins were extracted and desalted with S-Trap^TM^ mini columns according to the manufacturer’s instructions (ProtiFi™). Proteins were digested with trypsin (V5111, Promega). Peptide concentrations were determined using Pierce^TM^ BCA Protein Assay Kit. 25 µg of each sample were labelled with TMTpro^TM^ 16plex Label Reagent Set (A44520, Thermo Scientific^TM^) and pooled. 100 µg of the pooled sample were fractionated on a high pH liquid chromatography (HPLC, Agilent 1100 Series). 54 fractions were collected and concatenated to 18 samples which were submitted for LC-MS/MS analysis on a Velos Pro Orbitrap Elite™ (Thermo Scientific^TM^) mass spectrometer coupled to an Easy nanoLC 1000 (Thermo Scientific^TM^) with a 200 cm µPAC^TM^ column (PharmaFluidics). Data were analysed using MSFragger in Fragpipe 16.0 [51] and the UniProt human reference proteome database downloaded on the 14^th^ of June 2021 (20,845 protein entries including 245 potential contaminants). We allowed for two missed cleavages, a fragment and a precursor mass tolerance of 20 ppm each. A mass shift of 304.207146 Da was set for the N-term and lysine (K) representing the tandem mass tags (TMT), a mass shift of 57.0214560 Da for cysteine (C) representing carbamidomethylations and of 40.010600 Da for N-term acetylation.

### Statistical Analysis

Mass spectrometry data were analyzed using partial least-squares discriminant analysis (PLS-DA) and/or linear models of microarray analysis (LIMMA) to identify differentially abundant proteins. Those analyses were performed in the R environment [52] using the mixOmics [53] and limma [54] packages. Gene Ontology pathway analysis was performed using the topGO algorithm [55] in R. The results were visualised using the basic R plot functions, ggplot2 [56] and the EnhancedVolcano package [57]. In vitro cell based assays were analysed and visualised in Graphpad Prism 6. To identify significant differences ANOVA followed by either Sidak’s or Tukey’s test was applied.

### TP53 Functional Assay

Nuclear extracts from 5*10^6 cells were collected using a Nuclear Extraction Kit (Abcam, ab113747) according to the manufacturer’s instructions. The concentrations of the nuclear extracts were measured with the Pierce™ BCA Protein Assay Kit Reducing Agent Compatible (Thermo Scientific™, 23250). 10 µg of nuclear extract were used for the TP53 functional assay that was conducted according to the manufacturer’s instructions with the calorimetric p53 Transcription Factor Assay Kit (Abcam, ab207225). In short, nuclear extracts were incubated with a TP53 specific double stranded DNA sequence binding site immobilised on an ELISA plate. The bound TP53 was then detected with a TP53 antibody and a HRP conjugate whose OD signal was measured at 450 nm. The OD was normalised to 1 µg nuclear extract.

### Neocarzinostatin Treatment and Immunocytochemistry

Cells were seeded on coverslips (5000/coverslip, triplicates) coated with Poly-D-Lysine (Gibco^TM^) and grown overnight. Cells were treated with 250 ng/ml NCS for 1 hour and washed twice with DPBS. Fresh DMEM was applied and cells were allowed to recover for 1.5 hours. Cells were washed and fixed for 15 min with formalin. Fixed cells were permeabilized with 0.1 % Triton X-100 5 % BSA in DPBS, blocked with 5 % BSA in DPBS and incubated with the 1:1,000 diluted primary antibody for 2 hours. The secondary anti-rabbit AF555 coupled antibody or anti-mouse AF647 coupled antibody was diluted 1:10,000 and incubated for 1.5 hours. Cell nuclei were stained with Hoechst 33342 (B2261, Sigma-Aldrich). Coverslips were mounted (Prolong™ Gold antifade reagent, P10144 Invitrogen™, Thermo Fisher) on glass slides. The AF555, AF647 and Hoechst stains were inspected by microscopy. For the ɣH2A.X response calculations, the AF555 signal intensities and nuclei sizes were calculated using ImageJ. We calculated the ɣH2A.X response as the AF555 signal intensity per nucleus size of one cell.

### Comet Assay

Cells were seeded (4*10^5^/well) in 6-well plates, grown overnight and exposed to 500 ng/ml NCS for 1 hour. As positive control, cells were treated with 50 µM tert-Butyl hydroperoxide (B2633, Sigma-Aldrich). Cells were harvested, mixed 1:10 with 0.5 % low melting agarose (Biozym) and plated on microscopic slides, pre-coated with 1 % regular agarose (Biozym). Slides were incubated in cold lysis buffer (2.5 M NaCl, 100 mM EDTA, 1 % Triton X-100 and 10 mM Tris pH 10) at 4 °C for 1 hour. Lysis buffer was removed and exchanged to cold alkaline electrophoresis buffer (300 mM NaOH, 1 mM EDTA pH 13). After a 25 min incubation at 4 °C, the electrophoresis was conducted at 1 V/cm (22 V) at 4 °C for 15 min. The slides were washed with water, fixed with 70 % EtOH and air-dried. The DNA was stained with Vista Green DNA staining solution (238554, abcam) and DNA comets were inspected by microscopy. The DNA comets were analyzed using the OpenComet plugin [58] for imageJ. The olive moment was used to compare the conditions. Significant differences between groups were identified with the Kruskal-Wallis rank sum test in R. Pairwise comparisons were investigated performing the Wilcoxon rank sum test and p values were corrected using the Bonferroni method.

## Results and Discussion

### Global Proteome Profiling of iGBM

As an initial attempt to establish a proteomic platform for GBM (normal brain vs. iGBM), we performed explorative, mass spectrometry-based proteome profiling of cryo-preserved tissue specimens from five iGBM cases and five control samples of non-malignant brain tissue resected to approach the tumor site during surgery (Table 1). Non-patient matched pairs of one iGBM and one non-malignant brain sample were arranged (Fig. 1.a) and those pairs were labelled with stable isotope light and heavy formaldehyde. More than 4500 proteins were identified and quantified (Fig. 1b). Supervised partial least-squares discriminant analysis (PLS-DA) clearly separates the iGBM and non-malignant samples indicating distinct proteome compositions (Fig. 1c). Linear models of microarray analysis (LIMMA) were used to identify differentially abundant proteins (Fig. 1d). Based on this, a total of 588 dysregulated proteins (p_adjusted_-<0.05) were further used in a topGO enrichment analysis (215 upregulated and 372 downregulated proteins): In iGBM, upregulated biological processes include platelet degranulation, extracellular matrix organization, blood coagulation and fibrin clot formation as well as several immune response pathways (Fig. 1e). Among the downregulated biological processes dominate the mitochondrial ATP synthesis coupled electron transport as well as synaptic transmission and neurotransmitter pathways (Fig. 1f).

**Figure 1.**
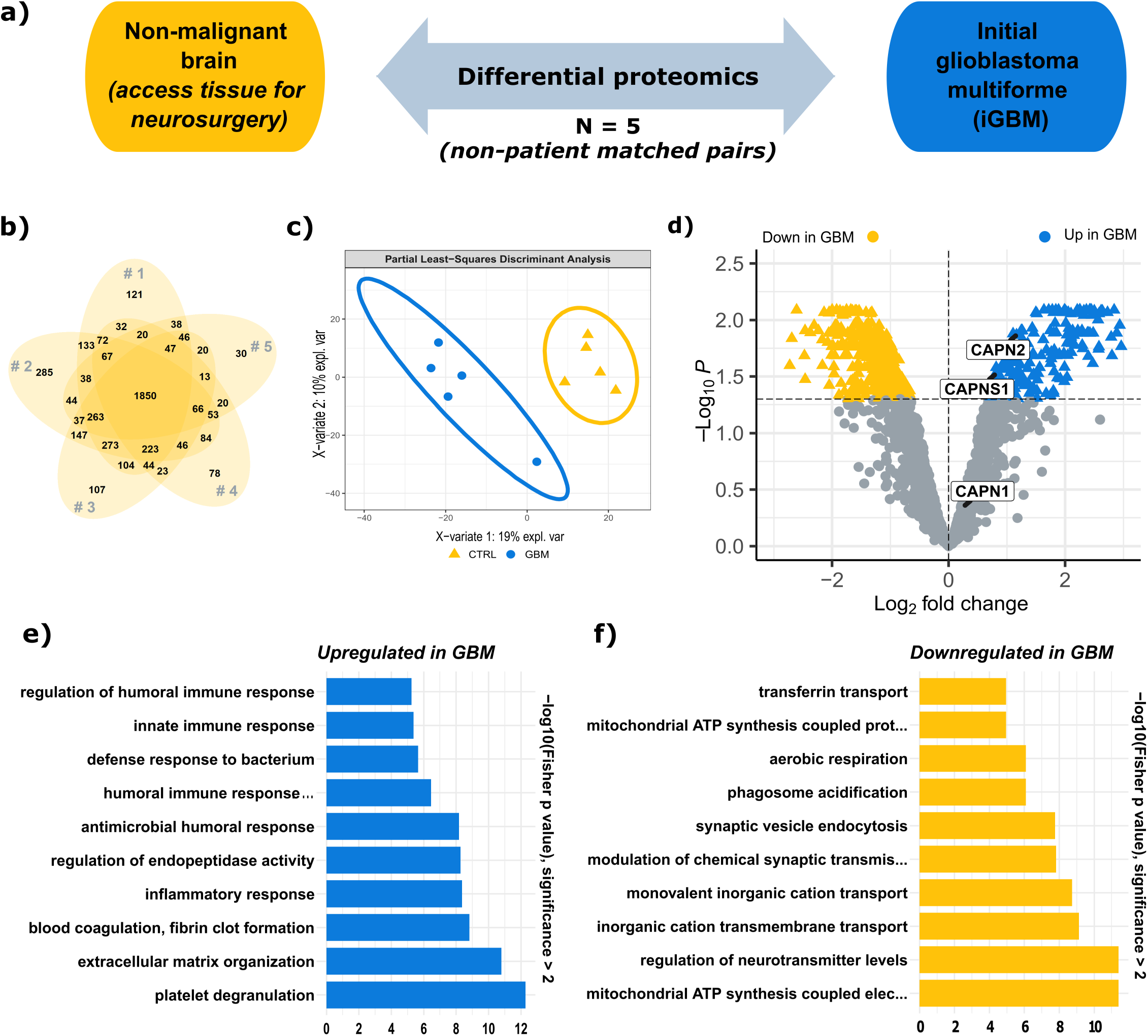
Proteome comparison of initial GBM and non-malignant brain (n=5). a) Schematic of the proteomic comparison. One iGBM sample was paired with one non-malignant brain sample. b) Venn diagram showing the overlap of all identified and quantified proteins in each of the five non-patient matched pairs. c) Partial least-squares discriminant analysis (PLS-DA) of the initial glioblastoma (GBM, blue dot) and control non-malignant brain (CTRL, yellow triangle) samples showing a clear separation. d) Volcano plot presenting the upregulated (blue triangle) and downregulated (yellow triangle) proteins in iGBM (LIMMA). e) Gene Ontology annotated biological processes upregulated in iGBM. f) Gene Ontology annotated biological processes downregulated in iGBM.

### Overexpression of Calpain-2 and the Calpain Small Subunit in glioblastoma

Detailed inspection of the upregulated proteins led us to notice overexpression of calpain-2 and the calpain small subunit in GBM (Fig. 1d). Interestingly, calpain proteases have only been marginally studied in the context of GBM: Although calpain proteases are recognized as important actors in cell biology with more than 10,000 PubMed entries, there are less than 60 entries for the search term “calpain + glioblastoma” as of 09/2022. Calpain proteases form heterodimers: They consist of a large catalytic subunit and a shared small regulatory subunit (CAPNS1). The predominantly expressed catalytic subunits are either calpain-1 (CAPN1) or calpain-2 (CAPN2). Our proteomic data shows that the protein abundance levels of calpain-2 and CAPNS1 are significantly upregulated in the iGBM samples with Log2 fold changes of 1.166 (p_adjusted_ <0.02) and 0.804 (p_adjusted_ =0.03), respectively (Fig. 2a). Calpain-1 is only slightly increased in iGBM with a Log2 fold change of 0.27 and a p_adjusted_ value of 0.44. Transcriptomic expression data obtained from the Cancer Genome Atlas (TCGA; www.cancer.gov) and GTEX (www.gepia2.cancer-pku.cn) show a general tendency for increased calpain expression in GBM tissue compared to non-malignant brains (Figs 2b-d). Calpain-2 expression levels are significantly increased (p <0.05), while calpain-1 and CAPNS1 show tendencies to increased expression levels. Comparing the intra-group expression levels in GBM, a wide expression level range is covered with patients showing very high expression levels and others presenting rather low or similar to non-malignant brain levels. Calpain detection in whole protein extracts from GBM and non-malignant brain samples reveals varying but rather increased calpain-1 and calpain-2 protein expression levels in GBM samples (Fig. 2e). Here, we again observed patient-specific expression levels. Immunohistochemical stainings for calpain-1 and calpain-2 confirmed the protein expression in iGBM (Fig. 2f). Expression of calpain-1 and calpain-2 was largely restricted to tumor cells.

**Figure 2.**
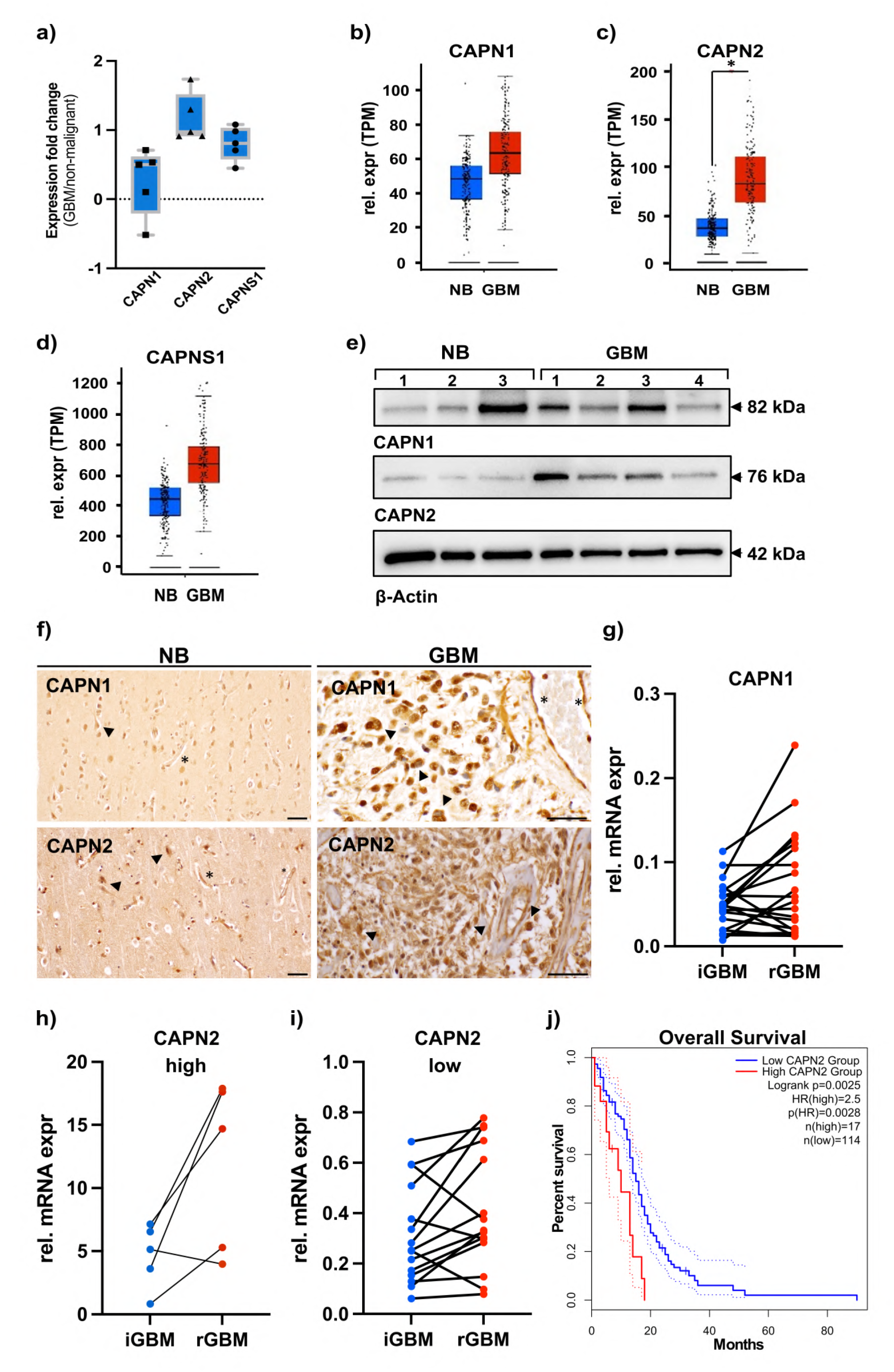
Higher abundance of calpains in initial and recurrent glioblastoma. a) Fc values for proteomic quantification of CAPN1, CAPN2, and CAPNS1 in five representative samples used for initial proteomic analysis shown in Fig. 1. b-d) expression of calpain-1 (CAPN1, b), calpain-2 (CAPN2, c), and calpain small subunit 1 (CAPNS1, d) in normal brain (NB) tissue and glioblastoma patient samples (GBM). Data were obtained from TCGA and GTEX data (www.gepia2.cancer-pku.cn) on n=207 (NB, blue bars) and n=163 (GBM, red bars) individuals. For CAPN2, expression difference is significant with p<0.01. e) Western Blot analysis of CAPN1 and CAPN2 from representative tissue samples of normal brain (access tissue, n=3) and tumor tissue (GBM, n=4). Molecular weights of CAPN1 and −2 are indicated by arrows on the right. As loading control, ß-actin was used. f) Immunohistochemistry for CAPN1 and CAPN2 in normal brain (NB, left) and sections from GBM (right) tissue. Neuronal staining is denoted by arrowheads in the left panel, and asterisks mark small blood vessels. In tumors, arrowheads denote tumor cells and asterisks large blood vessels with positive staining for CAPN1 and CAPN2 in endothelia. Scale bars in all images, 50 µm. g-i) expression of CAPN1 and CAPN2 in patient-matched initial GBM (iGBM) and recurrent GBM (rGBM) samples in 19 patients. g) CAPN1 expression is low and induced in 11 out of 19 patients. h) CAPN2 high expression in 5 out of 19 patients shows induction of CAPN2 in 4 out of 5 patient samples. i) CAPN2 low expression is induced in 11 out of 14 patient samples. In total, 4 patients showed a downregulation of CAPN2 in recurrent GBM, whereas 15 out of 19 show an increase in CAPN2 in recurrent tumors. j) Overall survival time of 131 patients with either high (red) or low (blue) calpain expression. Data were obtained from www.gepia2.cancer-pku.cn (accessed on 28^th^ September, 2022).

Increased abundance and activity levels of calpains have been reported for various human cancer types, while the implicated functions remain debatable [20–24]. In prostate cancer, over-expressed calpain degrades androgen receptors during chemotherapy-induced apoptosis, acting pro-apoptotic [59]. In contrast, calpain has also been reported to modulate androgen receptors leading to constitutive activity which eventually prevents therapy success [60]. In gliomas, the small regulatory CAPNS1 was proposed as a negative prognostic marker [20]. CAPNS1 was significantly upregulated in glioma patients and this correlated with worse overall patient survival. In the same study, the knockdown of CAPNS1 led to a damped migration ability suggesting involvement in focal adhesion and cell spreading. Nevertheless, there are multiple reports stating pro-apoptotic functions for calpain proteases under various conditions and in different cell types displayed by calpain inhibition preventing apoptosis [61–64]. Of note, most studies do not specify a calpain subunit, probably because the common inhibitors do not distinguish between them. With the present study, we aim to contribute to elucidating the role of upregulated calpain-2 in GBM.

### Recurrent GBM has a Tendency for Further CAPN2 Overexpression

Despite multimodal therapy for iGBM including surgical resection and radio-/chemotherapy, recurrence is common. We sought to investigate whether calpain expression levels are further increased in recurrent GBM (rGBM) as compared to iGBM. To analyze the mRNA expression levels of calpain-1 and calpain-2 in iGBM and rGBM, we performed qPCR on a second set of 19 matched patient samples with iGBM and rGBM (Table 2). Expression levels of calpain-1 were low in initial GBM and increased in 11 out of the 19 patients in recurrent GBM (Fig. 2g). For calpain-2, expression levels were variable with five patients showing high and 14 patients showing low expression (Figs 2h-i). 15 patients showed an induction of the calpain-2 expression in the recurrent GBM, whereas the other four patients showed a decreased expression in recurrent GBM. We conclude that recurrent GBM exerts a tendency for further calpain-2 overexpression. This finding implies that calpain-2 could contribute to therapy resistance and overall survival, in particular in those patients from our cohort with high initial calpain- 2 expression levels (patients 5,17,18, see Table 2). When the same distribution of calpain-2 expression levels was simulated in the TCGA patient cohort (20 % high vs. 80 % low), we found a strong correlation of high calpain-2 levels with significantly reduced survival (Fig 2j) and a Hazard ratio of 2.5 (p = 0.0028) suggesting that a high calpain-2 expression in tumors is of high relevance for patient survival. These findings prompted us to investigate the mechanistic role of calpain-2 in GBM therapy resistance.

### Calpains Contribute to Temozolomide Resistance in Primary GBM cells

Based on the previous observations, we hypothesized that calpains may contribute to the widely reported temozolomide (TMZ) resistance [5]. To test this, we treated three patient-derived GBM cells (Table 3) with the first-line chemotherapeutic TMZ and the synthetic calpain inhibitor PD150606 (PD). PD is a selective inhibitor for calpain proteases but does not distinguish between calpain-1 and calpain-2 [65]. Compared to TMZ administration alone, the combined administration of TMZ plus PD150606 led to a decreased viability measured with an MTT reduction assay (Fig. 3a).

**Figure 3.**
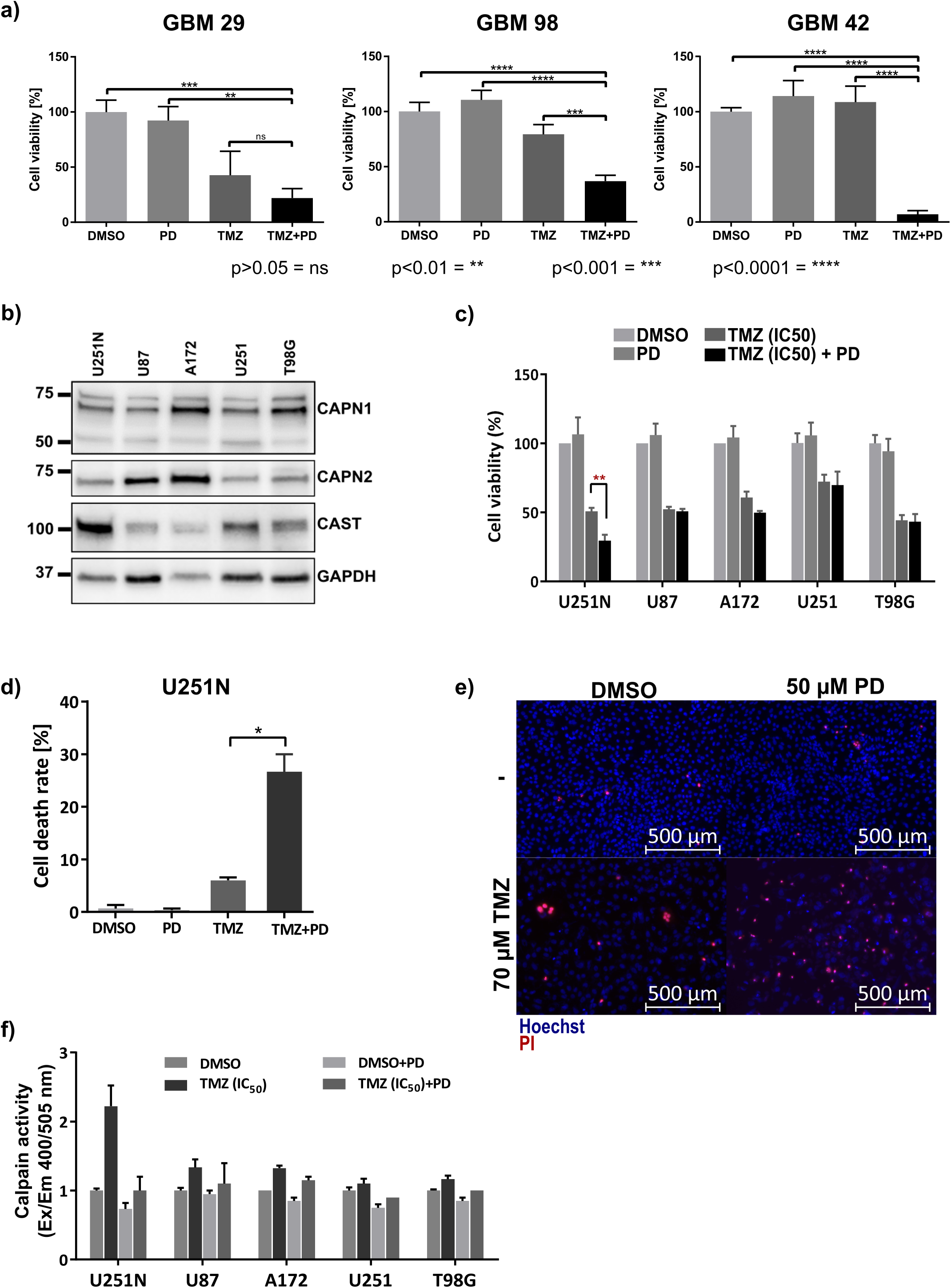
Temozolomide (TMZ) sensitivity of primary and established GBM cell lines. Error bars indicate the standard deviation. a) Cell viability (MTT reduction assay; ANOVA and Tukey’s test) of primary, patient-derived GBM cells after 5-day exposure to TMZ +/- calpain inhibition by PD150606 (PD). b) Western blot analysis of five selected GBM cell lines for their CAPN1, CAPN2, and calpastatin (CAST) expression. GAPDH was used as loading control. c) Cell viability (CellTiter Glo assay; ANOVA and Tukey’s test) of established GBM cell lines (n=3) after 5-day TMZ +/- PD administration. d) Cell death rate in U251N cells after 5-day TMZ (f.c. 70 µM) and PD (f.c. 50 µM) treatment. Cell death rate was counted as the number of propidium iodide (PI) positive cells relative to the total number of cell nuclei stained with Hoechst (n=3, ANOVA and Tukey’s test). e) Exemplary fluorescent images showing Hoechst stained nuclei (blue) and PI stained dead cells (red) after the indicated treatment. f) Calpain activity is increased in U251N cells upon 5-day TMZ administration (n=3). Calpain activity (fluorometric intensity (Ex/EM 400/505 nm)) is calculated relative to the DMSO treated control cells.

**Table 3:** Mutational statuses of primary patient derived and established GBM cell lines.

| GBM cell line | IDH1 R132H | TP53 | EGFR v III | MGMT status |
| --- | --- | --- | --- | --- |
| GBM 29 | negative | mutated | negative | unmethylated |
| GBM 98 | negative | mutated | negative | unmethylated |
| GBM 42 | negative | mutated | negative | unmethylated |
| U251 | negative | mutated | negative | methylated |
| U251N | negative | mutated | not tested | methylated |
| U87 | negative | wild type | negative | unmethylated |
| A172 | negative | wild type | negative | methylated [111] |
| T98G | negative | mutated | negative | unmethylated [111] |

Calpain inhibition via PD150606 enhanced TMZ sensitivity of primary GBM cells. Administration of TMZ alone resulted in a decreased viability of two primary cell lines (GBM 98: 1.3 times decreased; GBM 29: 2.3 times decreased; both p_adjusted_ <0.05). Compared to the sole TMZ exposure, additional inhibition via PD resulted in a synergistic effect: the viability of GBM 98 and GBM 29 was further lowered by 2.2 and 1.9 times (p_adjusted_ <0.01), respectively. In GBM 42 cells, viability was even 14.6 times reduced compared to the equimolar DMSO control, though TMZ alone was insufficient in inducing cell death. These results confirm our hypothesis that calpains contribute to the ability of GBM cells to withstand TMZ therapy.

### Calpains Contribute to Temozolomide Resistance in Selected Established GBM Cell Lines

Next, we aimed to confirm calpain-1 and calpain-2 expression in five established GBM cell lines: U251N, U251, A172, U87 and T98G (Table 3). All five cell lines express detectable levels of calpain-1 and calpain-2 (Fig. 3b). Additionally, we tested whether the cells express the endogenous calpain inhibitor, calpastatin (CAST). Indeed, all five cell lines express calpastatin and we observed the highest calpastatin level in U251N.

To investigate whether the established cell lines are sensitive to TMZ and whether they display the synergistic effect of TMZ treatment and calpain-1 and calpain-2 inhibition, we measured the cell viability upon TMZ and PD exposure with an ATP consumption assay (CellTiter-Glo® assay). All five cell lines were sensitive to TMZ and showed a diminished viability after five-day TMZ exposure (Fig. 3c). Except for U251N, the addition of the inhibitor did not further decrease viability. Notably, U251N viability was significantly reduced upon TMZ/PD co-administration compared to the TMZ alone (p_adjusted_ <0.01). We confirmed this observation by repeating this assay in U251N counting the rate of dead and alive cells with propidium iodide (PI) and Hoechst 33342 staining (Figs 3d-e). Concurrently, we observed an increased calpain activity under TMZ administration only in U251N cells (Fig. 3f). Enhanced calpain activation upon drug administration was already described in diverse pathological organs [66–72]. Cisplatin, a chemotherapy drug against germ cell tumors, leads to activation of calpains [73] while crosslinks with the urine bases in the DNA occur that lead to DNA damage and subsequent apoptosis induction [74], however no function has been ascribed to Calpains in this process.

Due to ongoing confusion around the U251 cell line and its subclones (including U251N), we performed Short Tandem Repeat (STR) profiling of the U251N cell line to confirm its authenticity (Table 4). The STR profiling identified the U251N used in this study as original U251N cells, as described by Torsvik *et al.*, 2014 [75].

**Table 4:** STR profiling of U251N cells to confirm cell authenticity.

| Locus | Chr. Loc. | ATCC marker | U251N used in study | Torsvik *et al.*, 2014 [75] |  |  |  |
| --- | --- | --- | --- | --- | --- | --- | --- |
|  |  |  |  | U251-MG | U-251-4q12 | U251N | U-251-FGA20gain |
| D3S1358 | Chr03 |  | 16/17 | 16/17 | 16/17 | 16/17 | 16/17 |
| TH01 | Chr01 | Yes | 9.3 |  |  |  |  |
| D21S11 | Chr21 |  | 29 | 29/30 | 29/30 | 29 | 29 |
| D18D51 | Chr18 |  | 13 | 13/15 | 13 | 13 | 13 |
| Penta E | Chr15 |  | 7/10 |  |  |  |  |
| D5S818 | Chr05 | Yes | 11/12 | 11/12 | 11/12 | 11 | 11 |
| D13S317 | Chr13 | Yes | 10/11 | 10/11 | 10/11 | 10/11 | 10/11 |
| D7S820 | Chr07 | Yes | 10/12 | 10/12 | 10/12 | 10/12 | 10/12 |
| D16S539 | Chr16 | Yes | 12 |  |  |  |  |
| CSF1PO | Chr05 | Yes | 11/12 |  |  |  |  |
| Penta D | Chr21 |  | 12 |  |  |  |  |
| AMEL | X/Y | Yes | X | X/Y | X/Y | X | X |
| vWA | Chr12 | Yes | 16/18 | 16/18 | 16/18 | 16/18 | 16/18 |
| D8S1179 | Chr08 |  | 13/15 | 13/15 | 13/15 | 13/15 | 13/15 |
| TPOX | Chr02 | Yes | 8 |  |  |  |  |
| FGA | Chr04 |  | 21/25 | 21/25 | 21/25 | 21/25 | 21/25 |

### Knockdown of Calpain-1 or −2 Sensitizes U251N Cells to Temozolomide Treatment

Following these observations, we knocked down CAPN1 (shCAPN1) or CAPN2 (shCAPN2) expression in U251N cells to corroborate the inhibitor-based experiments and to focus on individual calpain proteases. The constitutive, stable, polyclonal knockdowns reduced expression of either calpain-1 or calpain-2 by > 90 % as compared to a non-targeted control shRNA (termed shCTRL) (Fig. 4a). We employed two distinct shRNAs for each calpain gene to mitigate potential off-target effects (termed #1 and #2). U251N cells were then exposed for five days to 70 µM TMZ and the rate of dead cells was measured by counting dead and alive cells with PI and Hoechst 33342 staining. Knocking down calpain-1 or calpain-2 was already sufficient to significantly increase the rate of dead cells upon TMZ administration (Figs 4b-c). We observed that the knockdown of calpain-2 led to a higher rate of dead cells after TMZ administration compared to the knockdown of calpain-1.

**Figure 4.**
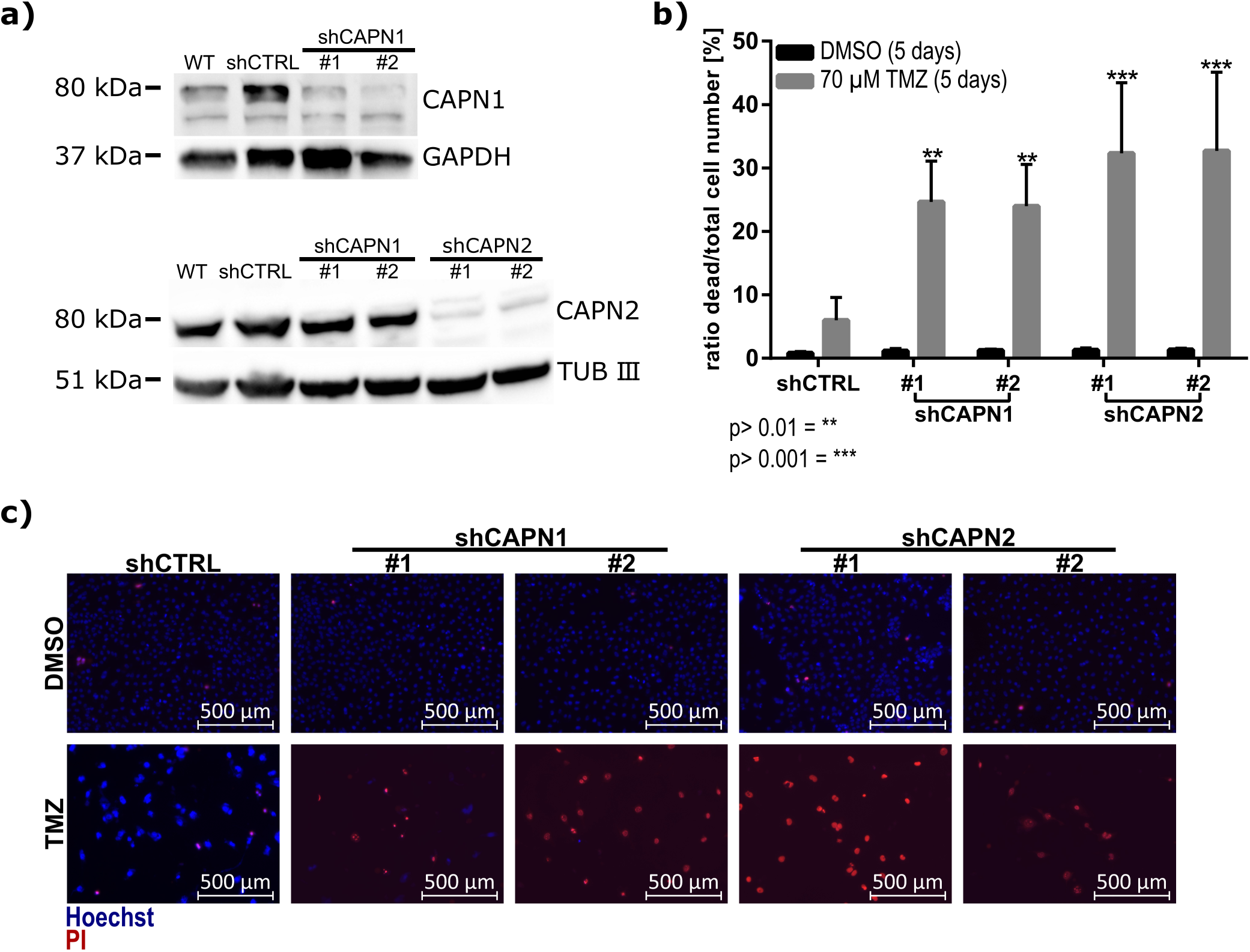
TMZ sensitivity of U251N cell deprived of calpain expression. a) Western blot analysis showing the shRNA expression knockdown of either CAPN1 or CAPN2 in U251N cells. We applied two shRNAs (termed #1 and #2) for CAPN1 and CAPN2, respectively. b) Cell viability of U251N deprived of either CAPN1 or CAPN2 compared to the shCTRL after 70 µM TMZ administration for 5 days (n=3, ANOVA and Sidak’s Test). Error bars indicate the standard deviation. c) Exemplary fluorescent images showing the propidium iodide (dead cells, red) and Hoechst (all cell nuclei, blue) stainings in U251N shRNA knockdowns.

After five days, both shCAPN2 replicates showed a higher rate of dead cells compared to the shCTRL: shCAPN2 #1 showed a mean 5.39-times higher cell death rate, shCAPN2 #2 a 5.44-times higher cell death rate. In addition, both shCAPN1 replicates also showed significantly increased cell death rates (shCAPN1 #1 =4.1-times higher, shCAPN1 #2 =4.0-times higher). Taken together, these results highlight a higher sensitivity against TMZ when hampering calpain proteases, especially calpain-2.

Next, we inhibited apoptosis by Q-VD-OPh, an irreversible inhibitor for caspase-3, −1, −8 and −9 [76, 77]. Q-VD-OPh effectively inhibited TMZ induced cell death (Fig. 5a). Necrostatin-1 (Nec-1), an inhibitor for necroptosis, failed to prevent cell death upon TMZ administration. This finding underlines that TMZ can induce apoptotic cell death. Upon TMZ administration increased levels of cleaved caspase-3 (CASP3), a key marker for apoptosis [78], were observed. Expression knockdown of either shCAPN1 or shCAPN2 led to further elevated levels of cleaved CASP3 (Fig. 5b). This observation raises the possibility that calpains are hampering the execution of TMZ-induced apoptosis. However, there is also the possibility that calpains rather act on an initial, pre-apoptotic level. Waterhouse *et al.*, 1998 [63] suggested that calpains are activated early after radiation-induced apoptosis and upstream of caspase activation.

**Figure 5.**
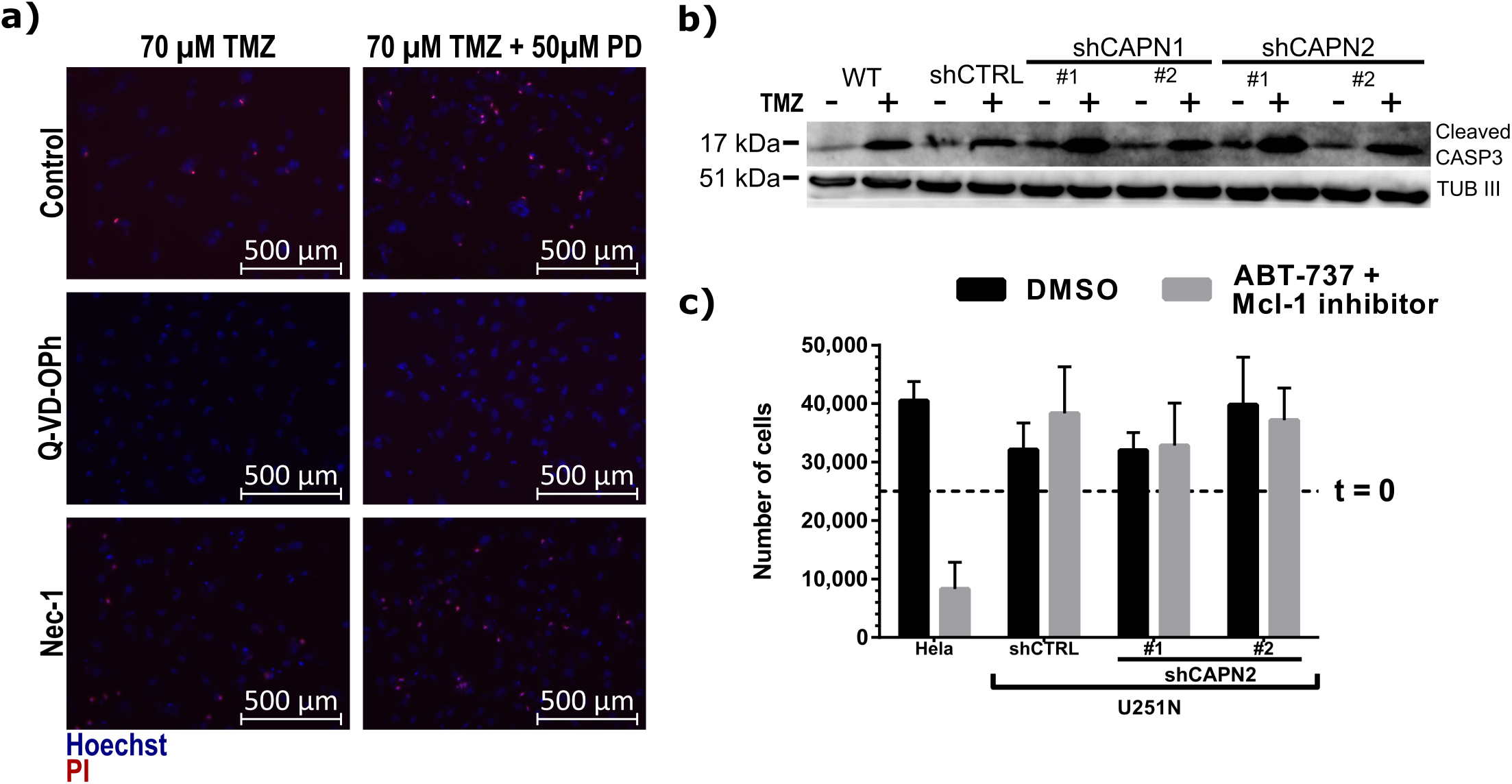
TMZ induced apoptotic cell death in U251N. a) Fluorescent images of U251N wild type cells after TMZ +/- PD administration. Apoptosis was inhibited with Q-VD-OPh, necroptosis with necrostatin-1 (Nec-1). Cell nuclei were stained with Hoechst (blue), apoptotic cells with propidium iodide (red). b) Western blot analysis of cleaved caspase-3 (CASP3) in U251N calpain knockdown cells +/- TMZ administration. b) Apoptosis induction in U251N cells. 2,5*10^4^ cells were seeded (t=0) and exposed to the BH3 mimetic ABT-737 and the Mcl-1 inhibitor. After 24 hours cells were stained with trypan blue (dead cells) and counted. Error bars show the standard deviation.

We sought to investigate this aspect by directly inducing apoptosis in U251N cells using the pro-apoptotic BH3 mimetic ABT-737 and Mcl-1 inhibitor [79]. However, both compounds failed to induce apoptosis in U251N cells (Fig. 5c).

### Proteome Comparison of U251N shCTRL vs. shCAPN2 Identifies Dysregulation of DNA Damage Sensing and DNA Damage Repair Proteins

From the previous results we concluded that calpain-2, when expressed in GBM cells, can protect cells against TMZ. To identify the underlying mechanism of calpain-2-dependent suppression of TMZ-induced apoptosis, we performed explorative, mass spectrometry based proteomics to compare treatment-naïve U251N shCTRL cells with both expression knockdown variants of calpain-2. We used five cell culture replicates (independently cultured and harvested) for each stable U251N knockdown line: 5x shCTRL, 5x shCAPN2 #1 and 5x shCAPN2 #2 in conjunction with TMT 16-plex to label and pool the samples. We extensively fractionated the pooled sample collecting 54 fractions, and concatenated those to 18 LC-MS/MS measurements. In total, 6044 proteins (including only unique peptides) were identified and quantified. We used LIMMA statistics to identify differentially abundant proteins between shCTRL and shCAPN2. Based on these data, 2,488 proteins were dysregulated (p_adjusted_ <0.05), demonstrating a massive proteome rearrangement solely mediated by calpain-2 (Fig. 6a). 1,243 proteins were significantly upregulated in cells deprived of calpain-2 expression. 1,245 proteins were significantly upregulated in the shCTRL cells.

**Figure 6.**
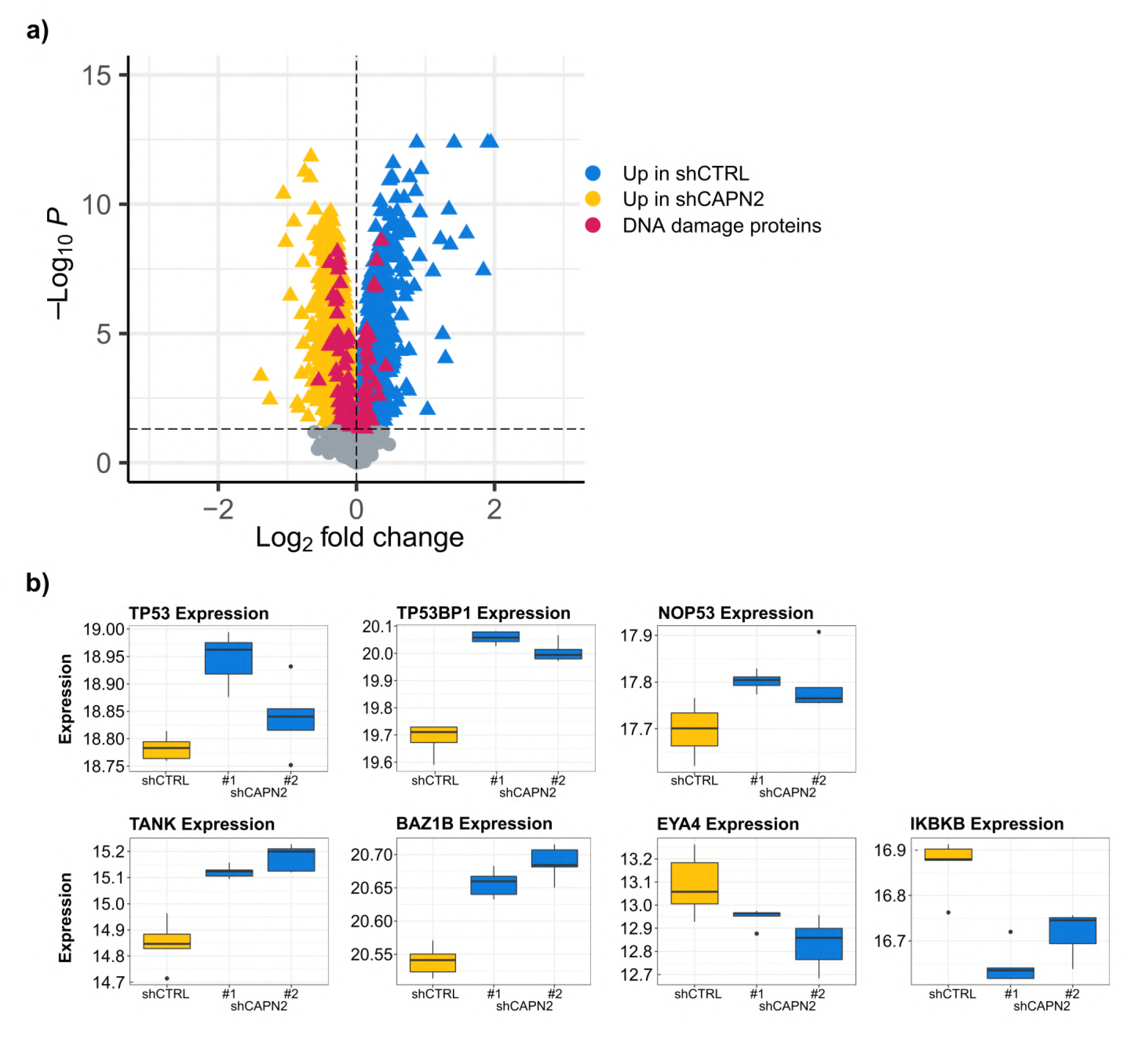
U251N proteome comparison of shCAPN2 and shCTRL cells (n=5). a) Volcano plot showing proteins upregulated (blue triangle) in shCTRL and upregulated (yellow triangle) in shCAPN2. DNA damage proteins are highlighted in pink. b) Expression of selected differentially expressed proteins (n=5). TMT reporter ion intensity normalized to median intensity of sample.

The proteome comparison with more than 6,000 identified proteins is probably the most complete portrayal on how calpain-2 affects proteome composition in glioblastoma cells. The high number of dysregulated proteins suggests a massive impact of calpain proteases in GBM cells. We identified the dysregulation of more than 100 proteins associated with DNA damage sensing and DNA damage repair (Figure 6a).

We found increased expression levels of TP53, TP53BP1 and NOP53 (Fig. 6b) in U251N cells deprived of calpain-2 expression. These proteins represent a cluster of tumor suppressing proteins [80–86]. TP53 is a sensor for DNA damage triggering DNA repair, but in case DNA repair is failing, its excessive nuclear accumulation induces apoptosis [87, 88]. TP53BP1 binds TP53 stabilizing the transcription of pro-apoptotic proteins and is rapidly recruited to sites of DNA double strand breaks (DSB), forming foci, and contributes to the DSB repair pathway choice [86,89–91].

Further, we detected upregulated expression levels of TANK and BAZ1B in shCAPN2 cells (Fig. 6b). TANK attenuates NF-κB activation upon DNA damage leading to cell death [92]. BAZ1B is a tyrosine kinase that phosphorylates ‘Tyr-142’ of histone H2AX [93, 94]. This phosphorylation is a key regulator of the DNA damage response critical for the cell’s fate as it impedes the recruitment of DNA repair complexes to phosphorylated ‘Ser-139’ of histone H2AX and promotes recruitment of pro-apoptotic factors. In contrast, we found EYA4 and IKBKB upregulated in shCTRL cells. The EYA protein family is BAZ1B’s natural antagonist as it dephosphorylates ‘Tyr-142’ of histone H2AX allowing for the recruitment of DNA repair factors [93]. IKBKB phosphorylates the inhibitor of NF-κB leading to NF-κB activation and improved cell survival [95–97].

Conclusively, these observations show the huge impact of calpain-2 expression on the U251N cells. Its presence/absence strongly affects the expression of DNA damage sensing and repair proteins. We hypothesize that expression of calpain-2 induces a “priming” of U251N cells to favor DNA repair over apoptosis when cells are exposed to DNA damage. The increased abundance of TP53 and TP53BP1 in calpain-2 silenced cells suggests an increased ability to recognize DNA damage and induce apoptosis when cells lack calpain-2 expression.

### Calpain-2 Reduces Neocarzinostatin-induced DNA Damage in U251N Cells

In order to investigate the hypothesis, whether calpain-2 expression desensitizes U251N cells to DNA damage, we measured the DNA damage levels in response to neocarzinostatin (NCS, Figure 7a). NCS is an antitumor enediyne antibiotic [98, 99] that acts as a DNA damage-inducing agent. It is abstracting deoxyribose atoms causing DNA double strand breakage (DSB) with DNA damage marker such as phosphorylated (ɣ-)H2AX becoming apparent in under three hours. [100]. We assessed the ability to recognize DNA damage by measuring the ɣH2AX signal and performed the Comet assay to measure actual DNA damage[101] levels [101] of U251N cells exposed to NCS.

**Figure 7.**
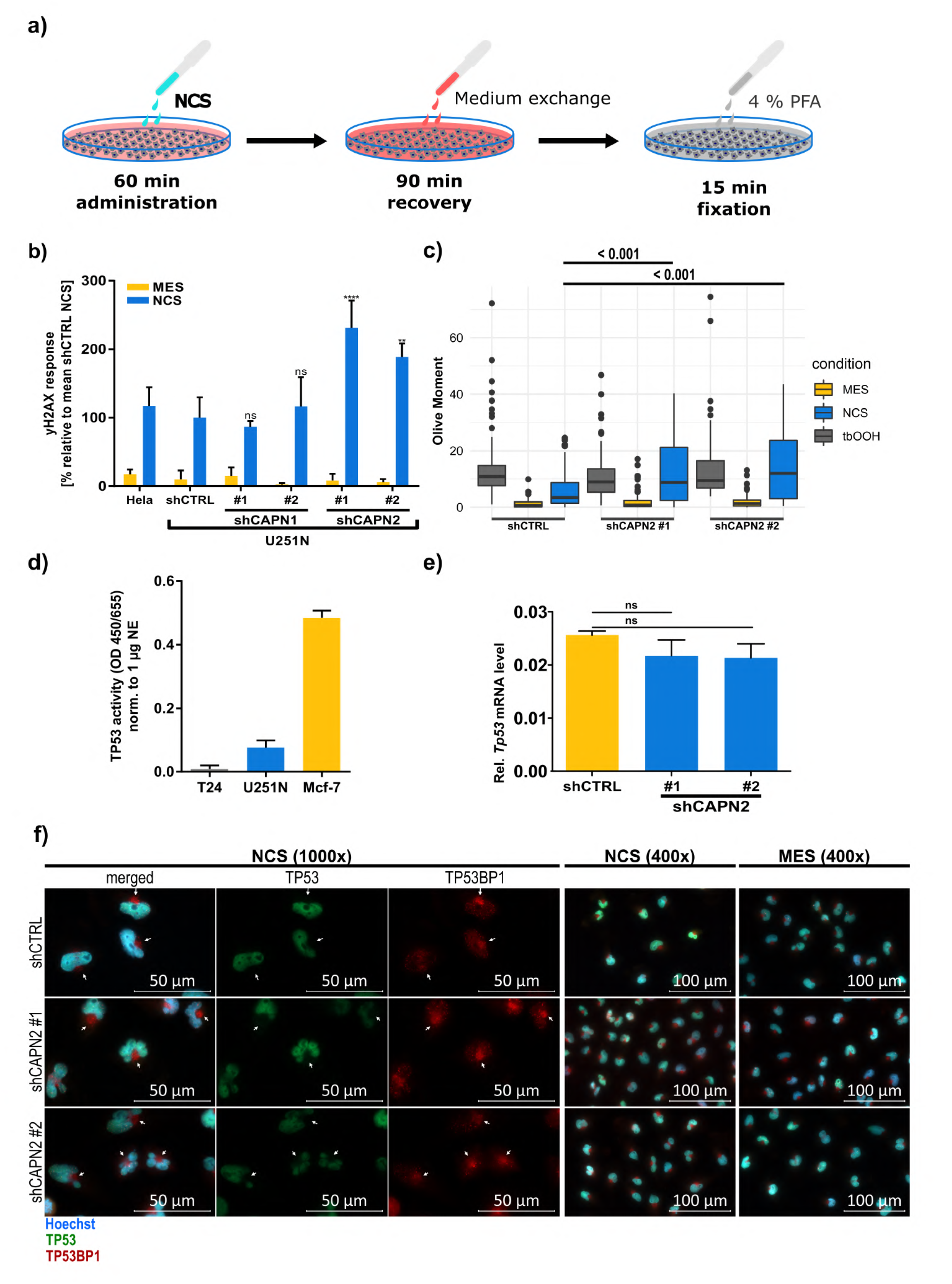
Neocarzinostatin (NCS) mediated DNA damage induction in U251N. a) Experimental procedure of the DNA damage induction. b) ɣH2A.X response of U251N and control Hela cells to NCS. U251N calpain knockdown cells were compared to the U251N shCTRL (ANOVA and Sidak’s test). Error bars indicate the standard deviation. c) Comet assay results assessing the DNA damage levels (olive moment) after NCS or tbOOH administration (Kruskall-Wallis ranksum test (p < 0.0001) followed by pairwise comparisons using the Wilcoxon rank sum test and Bonferroni correction). d) TP53 activity of U251N wild type cells in comparison to T24 (negative control) and Mcf-7 (positive control) cells normalized to 1 µg nuclear extract (NE). Error bars indicate the standard deviation. e) TP53 mRNA expression levels in U251N shCTRL and shCAPN2 knockdown cells indicate no difference. f) Fluorescent staining for TP53 (AF555, green) and TP53BP1 (AF647, red) in U251N shCTRL and shCAPN2 cells after NCS administration. Cell nuclei are stained with Hoechst (blue). Left panel shows 1000-x magnification with white arrows indicating TP53BP1 “hotspots” outside the nuclei. Right panel shows 400-x magnification of NCS treated and MES control cells.

The ɣH2AX signal was significantly less abundant in shCTRL cells (p < 0.05) (Figure 7b). Both shCAPN2 replicates showed significantly increased ɣH2AX signals by +131.5 % and +88.6 %, respectively. Phosphorylation of H2AX is an early event in the response to DSB, occurring within 1-3 minutes after DSB and leading to the formation of nuclear foci [102], where it attracts specifically repair factors such as MDC1 or TP53BP1 to DSB sites [102, 103].

The Comet assay showed significantly decreased DNA damage levels in shCTRL cells in response to NCS (Kruskal-Wallis rank sum test and Wilcoxon rank sum test p < 0.05; Figure 7c) which are 2.0 and 2.5 times decreased in comparison to shCAPN2 #1 and shCAPN2 #2, respectively. Interestingly, after exposure to tert-Butyl hydroperoxide (tbOOH), which induces single strand breaks (SSB) [104] and was used as a positive assay control, we did not observe any differences in the DNA damage levels.

This data proves that calpain-2 silenced U251N cells are more sensitive to recognize DSB, and suffer stronger DNA damage in response to DSB-induction. It also strengthens our hypothesis that calpain-2 (over-)expression yields an intrinsic “priming” of U251N cells to avoid DNA damage and apoptosis. Mechanistically, we suspect TP53 and TP53BP1 to play a pivotal role in the increased recognition of DNA damage upon calpain-2 silencing. Both proteins showed significantly increased expression levels and are known to be activated early after DSB.

### Calpain-2 Dependent Post-Translational TP53 regulation

TP53 is a known calpain substrate [28, 29], and calpain-dependent TP53 cleavage was shown to be necessary for G_1_/S-Phase transition upon DNA damage [105]. DNA damage induces TP53 activation, leading to apoptosis in order to clear cells with unstable genomes [83,106–108]. Increased nuclear TP53 levels were observed in Mcf-7 cells treated with calpain inhibitors, proving that the calpain inhibition stabilizes TP53 [28]. At the same time, truncated TP53 was hardly detectable *in vivo* suggesting that it is quickly degraded. To investigate the contribution of the increased TP53 levels in DNA damage sensing and repair when calpain-2 is silenced, we first examined, whether the mutated TP53 in U251N cells (Table 3) is functional. We performed an ELISA-based TP53 functional assay, which proved that TP53 retained a basal functionality (Figure 7d). U251N cells showed a lower TP53 activity than Mcf-7 cells (harboring wild type TP53) serving as a positive control, but a clear activity in comparison to the T24 cells, which are known to lack TP53 [109]. TP53 protein was exclusively localized in the cell nuclei, independent of the treatment (NCS or treatment-naïve; Figure 7f). The mRNA expression levels, measured via qPCR, did not differ between the control and calpain-2 silenced cells (Figure 7e). This observation suggests that TP53 is regulated post-translationally and not on a transcriptional level, which would be in line with reports about calpain-dependent TP53 cleavage. In the absence of calpain-2, TP53 shows increased protein levels in the cell nuclei. This can ultimately lead to the increased recognition of DNA damage and induction of apoptosis, which we observed in calpain-2 silenced cells. When calpain-2 is present, the U251N cells present decreased TP53 levels and impaired DNA damage sensing.

In addition, we found that, although significantly increased on the protein level, TP53BP1 transcription is not changed between the shCTRL and the first calpain-2 knockdown (shCAPN2 #1), but significantly reduced in the second calpain-2 knockdown (shCAPN2 #2; Figure 8a) We observed TP53BP1 in the cell nuclei, with specific accumulations in the nuclei that proved to be sites of DSB as co-localized with ɣH2AX (Figure 8b). We also observed “hotspots” of TP53BP1 expression very close to the nucleus but outside of it. TP53BP1 is rapidly recruited to sites of DSB and co-localizes with ɣH2AX [86,89,102,103]. It can bind TP53 to stabilize it and augment TP53-dependent transcription [90, 91].

**Figure 8.**
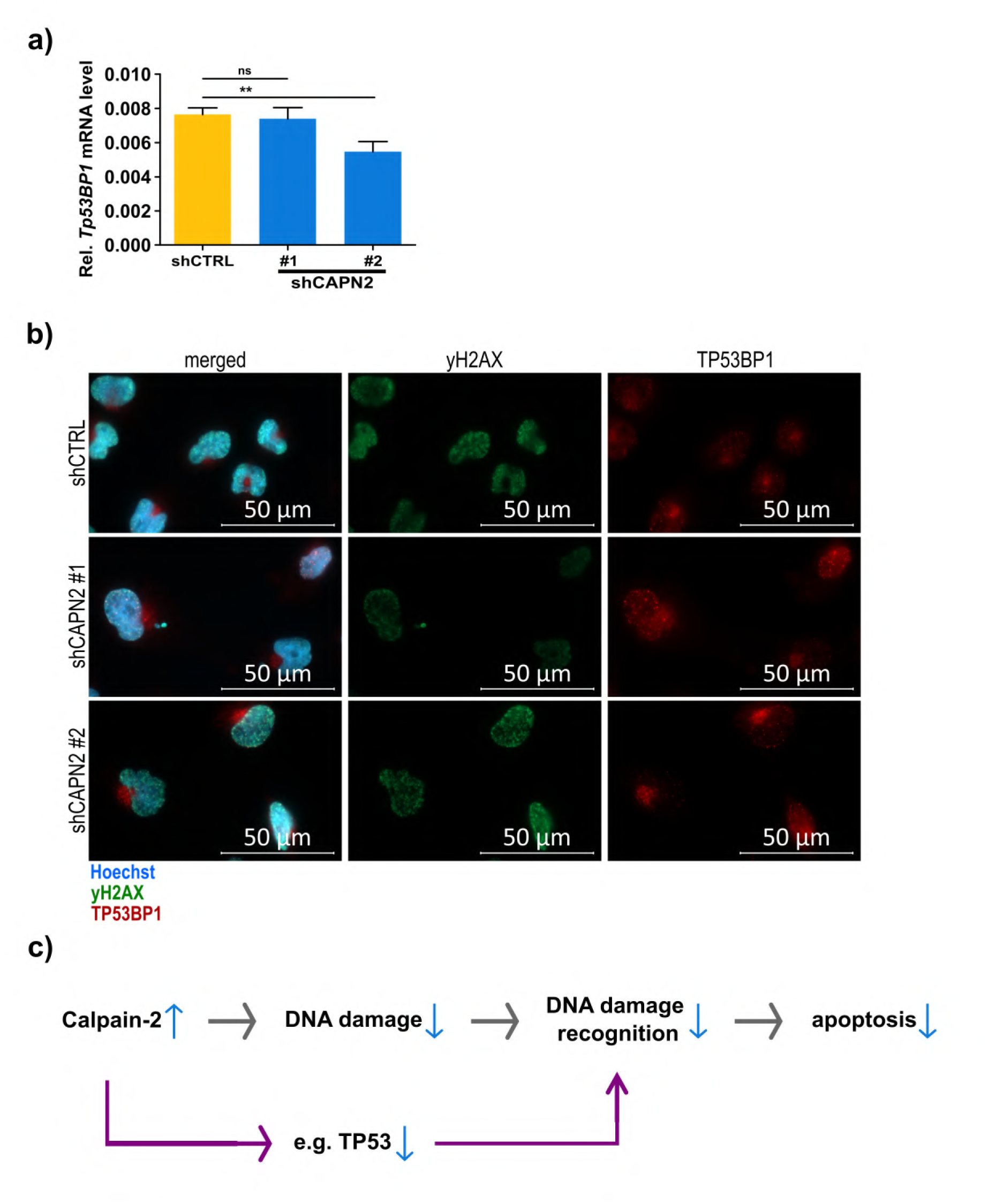
TP53BP1 expression and calpain-2 mediated DNA damage resistance mechanism. a) TP53BP1 mRNA expression levels in U251N shCTRL and shCAPN2 cells. b) Fluorescent staining for ɣH2AX (AF555, green) and TP53BP1 (AF647, red) in U251N shCTRL and shCAPN2 cells after NCS administration. Cell nuclei are stained with Hoechst (blue). c) Schematic of the calpain-2 driven DNA damage resistance mechanism. Decreased calpain-2 expression leads to a perturbed DNA double strand break repair, followed by increased DNA damage and an enhanced DNA damage recognition by e.g. TP53, and enhanced apoptosis induction.

Our data link calpain expression to TP53-dependent DNA damage recognition by observing a strong impact of calpain-2 expression on proteins associated with DNA damage sensing and DNA damage repair with more than 100 such proteins being significantly dysregulated upon calpain-2 silencing. Bearing in mind that the knockdown of calpain-2 expression augments the sensitivity to TMZ-induced apoptosis and enhances the recognition of DSB and levels of DNA damage upon NCS treatment, we assume that calpain-2 intervenes already at the level of DNA damage in the therapy resistance (Figure 8c). As TMZ alkylates DNA, leading to mismatch and DSB during replication [110]. The initial downregulation of TP53 and TP53BP1, the latter being especially important in DSB repair [91], by calpain-2 seems to prevent the U251N cells from DNA damage and subsequent apoptosis. Due to the very short NCS exposure time and the unchanged mRNA expression levels of TP53 and TP53BP1, we hypothesize that calpain-2 conveys an intrinsic “priming” to bypass DNA damage and compromise apoptosis. The rapid response of U251N cells upon NCS administration largely excludes a transcriptional and subsequent proteomic reprogramming of the cells to mitigate DNA damage. TP53, which was already described to be cleaved by calpain, seems to be the target protein inhibited by calpain to convey resistance. This model might be only applicable to this particular cell model, where TP53 expression is low and TP53 functionality is impaired but present at residual levels. However, the nuclear accumulation occurring under calpain inhibition might exceed a certain threshold of TP53-dependent DNA damage recognition to induce apoptosis.

## Conclusion

We conclude that calpain-2 contributes to TMZ resistance in GBM by impeding DNA damage recognition via the downregulation of DNA damage signaling proteins such as TP53 and TP53BP1. Calpain-2 and the small regulatory subunit CAPNS1 showed significantly increased expression levels in initial GBM as compared to non-malignant brain tissue. Calpain-2 showed further increased expression levels in recurrent GBM in a patient-specific manner. In addition, TMZ resistance in primary, patient-derived GBM cells and in established U251N cells was reduced by calpain inhibition. These observations show that calpain-2 contributes to the TMZ resistance and that it might be considered as a therapeutic target. However, since patients in our cohorts presented varying calpain-2 expression levels in initial and recurrent tumors, we suggest calpain-2 rather as a target for personalized care. We suggest that calpain-2 mediates a “priming” of GBM cells to prevent strong DNA damage and subsequent apoptosis. U251N cells with silenced calpain-2 expression showed increased sensitivity to NCS, which rapidly induces DNA double strand breakage similar to TMZ. The knockdown of calpain-2 led to altered patterns of DNA damage repair and response proteins. TP53 and TP53BP1 were significantly downregulated in U251N control cells. Calpain-2 silencing allowed for an increased abundance of these two proteins resulting in enhanced DNA double strand break recognition, and increased DNA damage levels. The calpain-2 knockdown rather favors apoptosis, whereas calpain-2 expression appears to promote DNA repair and cell survival. These findings have implications for the clinics: based on our results, GBM patients with high calpain-2 levels should be stratified and subjected to anti-calpain-2 treatment in combination with TMZ to achieve a tailored therapy and a better outcome.

## Abbreviations

GBM: glioblastoma
CAPN: calpain
CAPNS1: calpain small subunit 1
CAST: calpastatin
TMZ: temozolomide
NCS: neocarzinostatin
LC-MS/MS: liquid chromatography tandem mass spectrometry.

## Declarations

### Ethics approval and consent to participate

The study design was approved (Approval No. 185/11) by the Institutional Ethics Committees of the Medical Faculty of Marburg University. All participants in the study provided informed consent before specimens were collected.

### Consent for publication

All authors have agreed to publish this manuscript.

### Availability of data and material

The LC-MS/MS raw data from the proteomic comparison of U251N cells are available on MassIVE (ftp://MSV000089297@massive.ucsd.edu; password: a). The LC-MS/MS raw data from patients were not published as we consider proteomic patient data as sensitive, personal data. Upon request, these data can be made available upon submission of a data access agreement in accordance to the European standardization framework for data integration and data-driven in silico models for personalized medicine – EU-STANDS4PM. All other data are available upon request from the corresponding authors.

### Competing interests

There are no conflicts of interest to declare.

### Funding

This work was funded in part by Rhön Klinikum AG, grant number FI_35_2016 (J.W.B.), by the von-Behring-Röntgen Stiftung, grant number vBR 64-0018 (J.W.B.), and by the Federal Ministry of Education and Research (BMBF), ERANET PerMed Project “PerProGlio”, grant numbers 01KU1915B and 01KU1915 to J.W.B and O.S., respectively.

OS acknowledges funding by the Deutsche Forschungsgemeinschaft (DFG, projects 446058856, 466359513, 444936968, 405351425, 431336276, 43198400 (SFB 1453 “NephGen”), 441891347 (SFB 1479 “OncoEscape”), 423813989 (GRK 2606 “ProtPath”), 322977937 (GRK 2344 “MeInBio”), the ERA PerMed programme (BMBF, 01KU1916, 01KU1915A), the German-Israel Foundation (grant no. 1444), and the German Consortium for Translational Cancer Research (project Impro-Rec), funding by the MatrixCode research group, FRIAS, and support by the Fördergesellschaft Forschung Tumorbiologie, Freiburg im Breisgau.

### Authors’ contributions

OS and JWB conceived and supervised the study and provided funding; MNS, CC, ZWL, ML, and AS performed experiments, data analysis and data curation; AP and CN provided resources and clinical information; MNS wrote the manuscript draft; OS, JWB, and AP, edited the final manuscript. All authors read and approved the final manuscript.

## Acknowledgements

We want to thank R. Hannen for help with initial cell culture experiments as well as S. Motzny, and S. Stei for their excellent technical assistance. We would also like to thank R. Geiss-Friedlander and O. Bolgi for the support with the neocarzinostatin experiments as well as U. Maurer for supporting us with the apoptosis assays and for providing aliquots of ABT-737 and the Mcl-1 inhibitor.

## Supplementary Files

Additional File 1: IC_50_ curves for TMZ of five established GBM cell lines.

## Notes

ftp://

### Competing Interest Statement

The authors have declared no competing interest.

