## Supplementary figures and images for "Changes in calpain-2 expression during glioblastoma progression predisposes tumor cells to temozolomide resistance by minimizing DNA damage and p53-dependent apoptosis"

### Additional_File_1

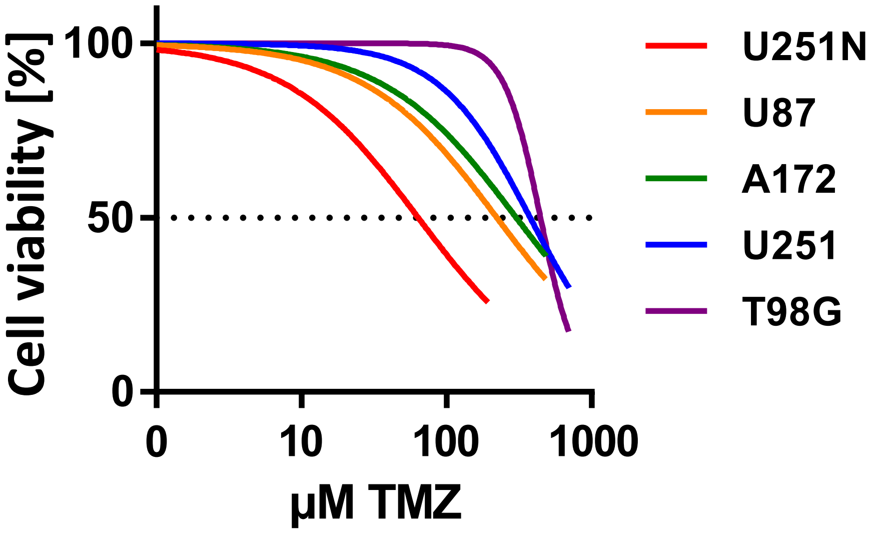
